# A dual role for GLI3 signaling in neural crest development

**DOI:** 10.1101/2025.08.31.673339

**Authors:** Simon J. Y. Han, Vinit Adani, Edward Farrow, Bhavalben Parmar, Ching-Fang Chang, Kim Cochran, Paige J. K. Ramkissoon, Ezekiel Esteban, Kelsey H. Elliott, Kevin A. Peterson, Brian Gebelein, Martín García-Castro, Samantha A. Brugmann

## Abstract

Neural crest cells (NCCs) are a multipotent cell population that undergo specification, epithelial-to-mesenchymal transition, migration, and differentiation into a plethora of cell types. A wealth of studies across various embryonic model systems have established dogma as to the molecular mechanisms and signaling cascades that contribute to NCC development. While Wnt, FGF, and BMP signaling pathways have well-established and essential roles in several aspects of NCC development, the Hedgehog (HH) signaling pathway has received limited attention for any specific role in this process. Herein, we propose two distinct, temporal roles for the transcription factor GLI3 in NCC development. *Gli3*, and other members of the HH pathway, were robustly co-expressed with established NCC induction and specification markers in chick, mouse, and human embryonic stem cell derived NCCs. Early knock-down of GLI3 reduced expression of key markers of NCC specification and conditional knock-out of *Gli3* post-specification specifically impaired the ability of cranial NCCs to differentiate into ectomesenchymal derivatives. Together, these results demonstrate dual, novel roles for GLI3 in early NCC specification and later in cranial NCC differentiation.

**Summary Statement:** GLI3 plays distinct temporal roles in neural crest specification and differentiation into ectomesenchymal derivatives.

## Introduction

Neural crest cells (NCCs) are a transient, multipotent embryonic population of cells derived from the ectoderm. NCCs are induced early in embryonic development, specified (Stuhlmiller and Garcia-Castro, 2012, de Croze et al., 2011, Selleck et al., 1998, Prasad et al., 2012), undergo an epithelial-to-mesenchymal transition, and migrate to distal targets throughout the body where they differentiate into a multitude of cell types (Leathers and Rogers, 2022, Zhang et al., 2014, Kerosuo and Bronner-Fraser, 2012, Bronner and LeDouarin, 2012). NCCs along the axial level of the embryo (cranial, vagal, cardiac, trunk, and sacral) are endowed with differing levels of developmental potential, with cranial neural crest cells (CNCCs) having the broadest degree of potency. CNCCs make robust contributions to numerous head structures, where they differentiate into both non-ectomesenchymal (e.g., neurons, glia, pigment cells) and ectomesenchymal (e.g., bone, cartilage, and smooth muscle) derivatives (Perera and Kerosuo, 2021, Noden, 1983, Weston et al., 2004). This exceptional ability has been of particular interest in the field (Moore Zajic et al., 2025, Buitrago-Delgado et al., 2015, Zalc et al., 2021, Pajanoja et al., 2023).

Neural crest induction and specification has most commonly been thought to occur post-gastrulation, at the boundary between the neural plate and the non-neural ectoderm. At this junction, WNT, Bone Morphogenetic Protein (BMP), and Fibroblast Growth Factor (FGF) signaling initiate a transcriptional program (e.g., *Pax3, Pax7, Msx*, and *Zic1*) that defines the neural plate border (Garcia-Castro et al., 2002, LaBonne and Bronner-Fraser, 1999, Marchant et al., 1998, Mayor et al., 1997, Monsoro-Burq et al., 2003, Sauka-Spengler and Bronner-Fraser, 2008, Stuhlmiller and Garcia-Castro, 2012). Within this region, both NCCs and placodal cells are specified, however, NCCs express a unique combination of transcription factors including NCC specifiers such as *Tfap2a*, *Snail2, Sox9*, and *FoxD3* (Bronner and LeDouarin, 2012, Martik and Bronner, 2017), while the pre-placodal domain is more anterior and defined by the expression of other transcription factors including *Six1* and *Eya1* (Brugmann et al., 2004, Pandur and Moody, 2000, Almasoudi and Schlosser, 2021, Riddiford and Schlosser, 2017, Streit, 2007).

Despite this commonly accepted mechanism, more recent studies have suggested NCC induction occurs prior to neurulation (Basch et al., 2006, Prasad et al., 2020). A defined region of *PAX7+* epiblast cells contribute to the neural folds and migrating NCCs (Basch et al., 2006, Prasad et al., 2020, Betters et al., 2018, Patthey et al., 2008, Patthey et al., 2009). Independent of mesoderm or neural plate influence, loss of *PAX7* prevented the expression of several NCC markers including *SLUG*, *SOX9, SOX10* and *HNK-1* (Basch et al., 2006). Furthermore, a study in rabbit reported NCC specification occurred during gastrulation and was dependent on FGF signaling (Betters et al., 2018), and a recent report demonstrated that the nuclear receptor *Nr6a1* was required mid to late gastrulation for proper NCC specification in mouse embryos (Moore Zajic et al., 2025). Together, these studies suggest that NCC induction and specification may occur earlier than previously acknowledged, and that additional studies will be required to precisely determine when and how NCCs are induced.

While several key signaling pathways active during embryogenesis contribute to NCC development, the role of the Sonic hedgehog (SHH) pathway has not been thoroughly investigated in this process (Selleck et al., 1998). This lack of association is intriguing as the SHH pathway has a well-defined role in craniofacial patterning (Bush and Jiang, 2012, Chang et al., 2016, Millington et al., 2017), and mutations in ligands, receptors, and modulators of the pathway result in phenotypes that affect cranial NCC derivatives (Rowton et al., 2022, Ishikawa Y et al., 2021, Ishikawa et al., 2017, Yang et al., 2019, Elliott et al., 2018, Millington et al., 2017). The SHH pathway is mediated by bimodal GLI transcription factors (GLI TFs), which activate or repress target genes (Hui and Angers, 2011, Sasaki et al., 1999). GLI3, in particular, is an interesting candidate for playing a role in NCC development, as it is expressed in the dorsal neural tube, the site of NCC specification and epithelial to mesenchymal transition.

Herein we report two distinct, yet essential roles for GLI3 in both the induction/specification and differentiation of NCCs. *GLI3* is expressed during both NCC induction and specification across several species. Knockdown of *GLI3* prior to NCC specification results in a reduction of key NCC specifiers, such as *SOX10*, *FOXD3*, and *PAX7*. Furthermore, post cranial NCC specification, GLI3 binds at loci near genes involved in cranial NCC ectomesenchymal differentiation and conditional loss of *Gli* TFs in cranial NCCs results in impaired ectomesenchymal differentiation. Taken together, these data suggest GLI3 has both an early role in NCC specification and later role in cranial NCC differentiation.

## Results

### Hedgehog family members are expressed in developing neural crest cells

To determine if and how the HH pathway contributed to NCC development, we examined several publicly available datasets from various species. First, we profiled a *Tfap2aE1-eGFP+* FACS-enriched chick NCC RNA-seq dataset from various timepoints of NCC development (HH6-16) (Hovland et al., 2022) (**Fig. 1A**). Genes in the WNT, BMP, and FGF pathways involved in NCC induction (e.g., *CTNNB1*, *SMAD1*, and *MAPK3)* and specification (e.g., *TFAP2A, FOXD3, PAX7*) were highly expressed throughout NCC developmental stages. Interestingly, members of the HH signaling pathway were also expressed during various windows of NCC development at levels comparable to genes with well-established roles (**Fig. 1A**). Expression of *SHH* itself was detected in NCC populations, suggesting possible autocrine or paracrine signaling mechanisms in NCCs. Notably, expression of *GLI3* was enriched in NCCs compared to whole embryos (WE) (**Fig. 1A**). A second avian dataset which consisted of single-cell RNA sequencing (scRNA-seq) on early chick embryos (HH5, 1 somite, 4 somite, 7 somite) clustered by germ layer was also analyzed for NCC induction and specification genes (**Fig. S1A-S1B**). Analysis of HH signaling pathway genes confirmed *GLI3* expression in the ectoderm (**Fig. S1A, S1C**). Thus, these data suggested that members of the HH signaling pathway were expressed in the proper temporospatial domains to participate in NCC development.

**Figure 1.**
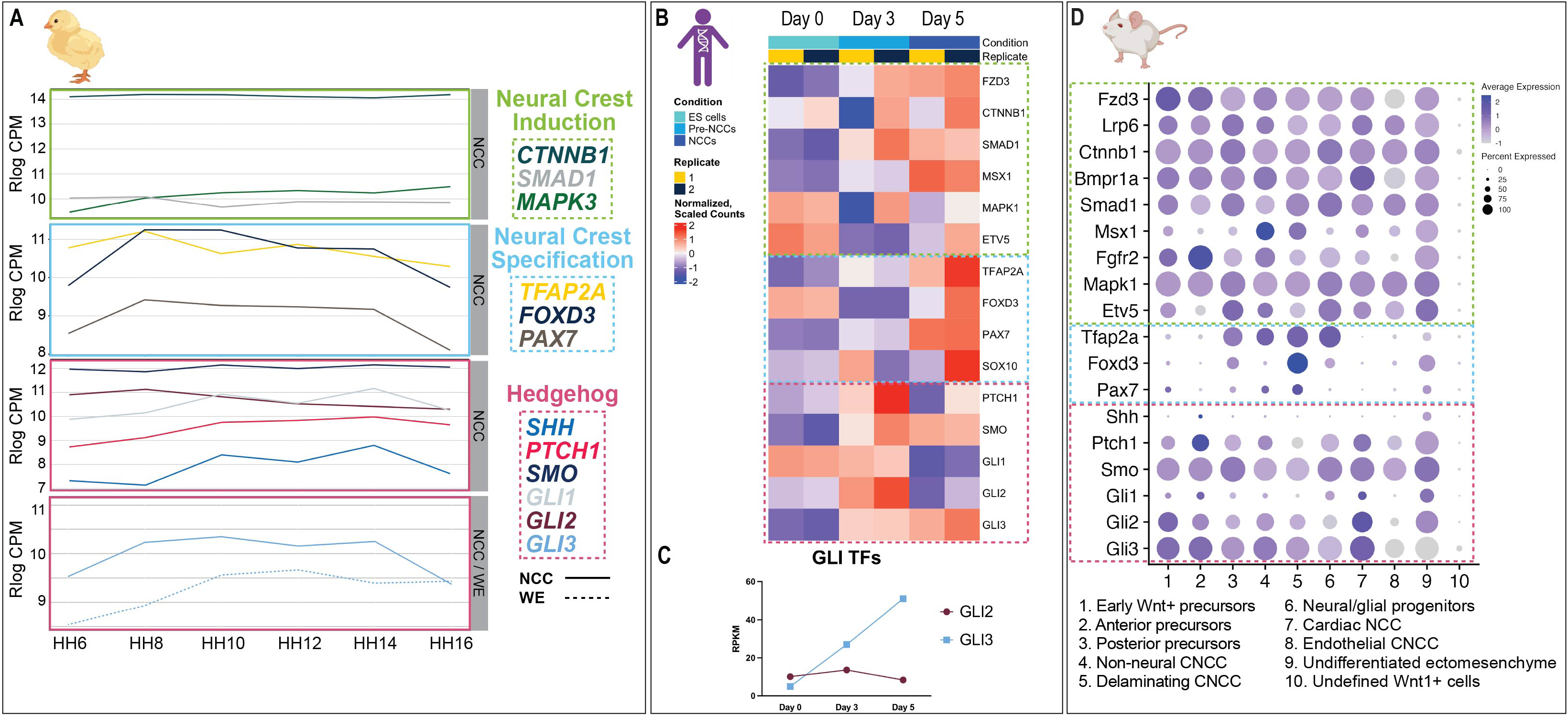
Hedgehog family members are expressed in developing neural crest cells. (A) Plots using an embryonic chick RNA-seq dataset showing expression of neural crest induction genes, neural crest specification genes, and HH pathway genes. *GLI3* expression is shown in both neural crest cells (NCC) and whole embryos (WE). (B) Heatmap of RNA-seq dataset from human embryonic stem cells during neural crest differentiation showing expression of neural crest induction genes, neural crest specification genes, and HH pathway genes. (C) Reads per kilobase million (RPKM) of GLI TFs from heatmap in (B). (D) DotPlot of a murine embryo scRNA-seq dataset reanalyzed to show expression of neural crest induction genes, neural crest specification genes, and HH pathway genes. Animal model schematics were created using Biorender.com.

While analysis of chick datasets clearly indicated expression of HH family members, expression in human samples had yet to be tested. To determine if HH family members were expressed during human NCC development, reanalysis of a deposited bulk RNA-sequencing experiment (Leung et al., 2019) performed on human embryonic stem (ES) cells during NCC differentiation (Day 0 (ES cells), Day 3 (Pre-NCCs), and Day 5 (NCCs)) was conducted. Several markers of NCC induction and specification were expressed at low levels in ES cells (Day 0). Expression of genes associated with NCC induction were increased in pre-NCCs (Day 3). By Day 5, when cells were fully differentiated into NCCs, expression of induction and specification markers was robust (**Fig. 1B**). Examination of HH family members revealed relatively low levels of expression at Day 0, a spike in expression of all family members in pre-NCCs (Day 3), and sustained expression of *SMO* and *GLI3* in NCCs (Day 5) (**Fig. 1B; Fig. S1D**). Of note, expression of *GLI3*, increased throughout NCC development while expression of *GLI2* remained consistent (**Fig. 1C**).

To further test for HH pathway expression in NCCs, we next analyzed a publicly available murine scRNA-seq dataset, which consisted of *Wnt1+* FACS-enriched cranial NCCs harvested between E8.0-E8.75 (4 somites to 10 somites) (Zalc et al., 2021) (**Fig. S1E**). Reanalysis of this dataset revealed expression of several known drivers of cranial NCC induction and specification. Members of the WNT (e.g., *Fzd3, Lrp6*, and *Ctnnb1*), BMP (e.g., *Bmpr1a*, *Smad1*, and *Msx1*), and FGF (e.g., *Fgfr2*, *Mapk1*, and *Etv5*) pathways involved in cranial NCC induction were robustly expressed in several NCC populations (**Fig. 1D, Fig. S1F**). Expression of key NCC specifiers (e.g., *Tfap2a, Foxd3* and *Pax7*) were also detected; however, expression was most robust in delaminating cranial NCCs (**Fig. 1D)**. Examination of HH family members revealed that *Shh, Ptch1, Smo, Gli1, Gli2, and Gli3* were expressed in most cranial NCC clusters at levels comparable to genes previously associated with NCC induction and specification, with the most robust expression in early neuroepithelial precursors, early *Wnt1+,* anterior/posterior, and non-neural cranial NCCs (**Fig. 1D**). Thus, analysis of several previously published datasets confirmed the expression of HH pathway members throughout the course of NCC development in various species. Of the HH family members examined, *Gli3* was of particular interest due to more robust expression in NCCs compared to whole embryo (**Fig. 1A**) and compared to *Gli2* (**Fig. 1B-D; Fig. S1A and S1C**). Thus, we continued our analysis with a focus on *Gli3*.

### GLI3 is expressed in developing neural crest cells

To further analyze the temporal and spatial expression of GLI3 during NCC development, both gene and protein expression were analyzed across species *in vivo*. *In situ* hybridization and RNAscope for *GLI3* in chick embryos across developmental stages (HH7-HH9) revealed *GLI3* expression in the headfold and dorsal neural tube (**Fig. 2A, Fig. S2A, S2B**). To validate GLI3 protein expression in NCCs, immunofluorescence (IF) was performed. IF at HH8 showed co-expression of GLI3 with the NCC marker PAX7 (**Fig. 2B**). To determine the expression of GLI3 in human NCCs, western blot analysis was performed using protein extracts from human ES cells and Day 5 NCCs. Expression of both full-length GLI3 (GLI3FL) and the truncated repressor (GLI3R) were negligible in ES cells. In contrast, both GLI3FL and GLI3R were robustly expressed in human NCCs (**Fig. 2C)**.

**Figure 2:**
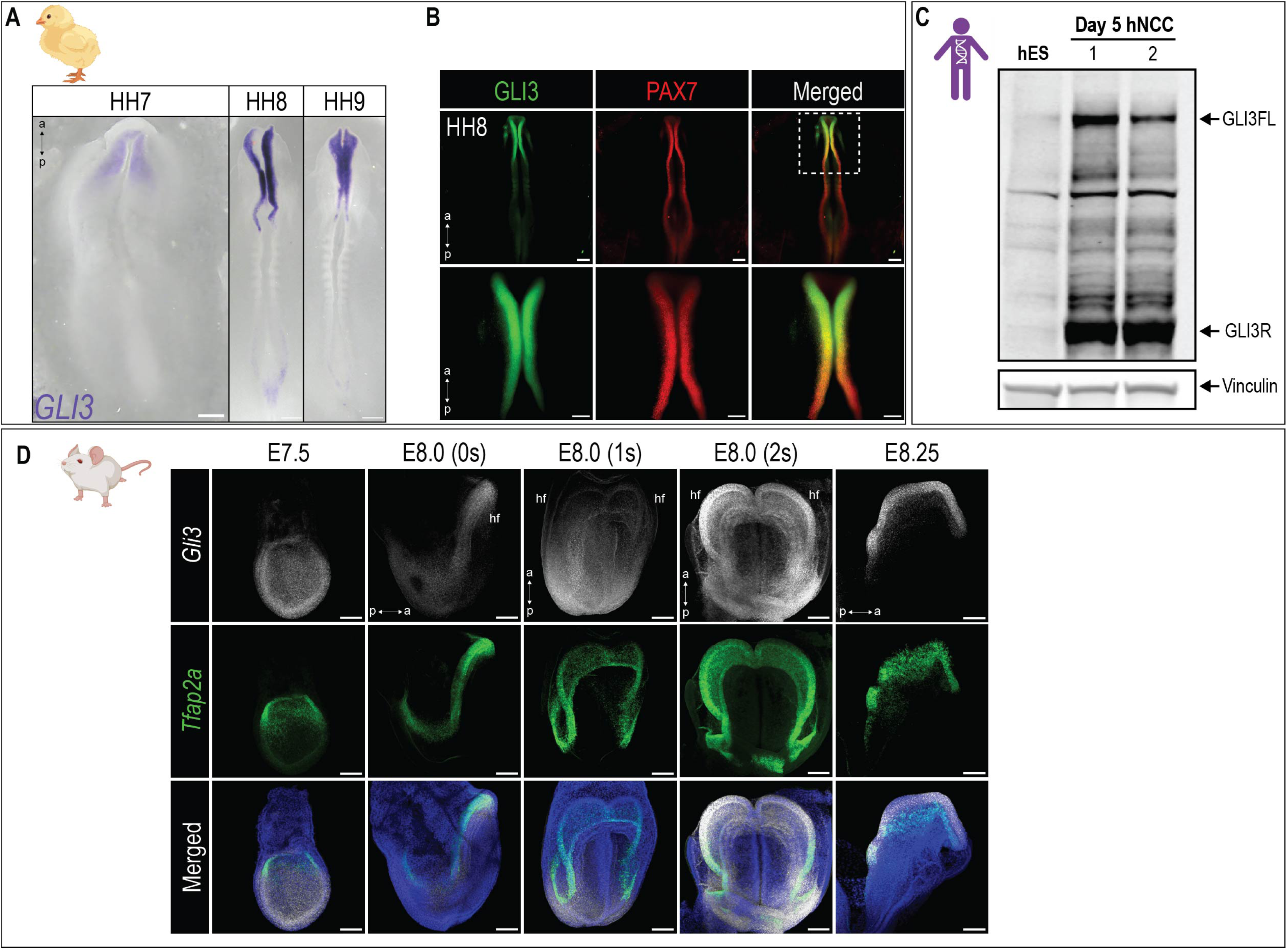
GLI3 is expressed in developing neural crest cells. (A) *In situ* hybridization for *GLI3* in chick embryos. (B) Immunofluorescence for GLI3 and PAX7 in HH8 chick embryos. (C) Western blots for GLI3 in ES (Day 0) and NCC (Day 5). (D) Whole-mount RNAscope of early mouse embryos showing co-expression of *Gli3* and *Tfap2a* in developing cranial NCCs. Animal model schematics were created using Biorender.com. (hf) headfold. Scale bars: 50μm (A, HH7); 100μm (A, HH8-9, B, D).

In murine embryos, *Gli3* transcripts were co-expressed with the NCC marker, *Tfap2a* from the onset of expression at E7.5 in the headfold. Co-expression of *Gli3* and *Tfap2a* was maintained in the headfold from the onset of *Tfap2a* expression at E7.5 through E8.25 (**Fig. 2D**). *Gli2* was also detected in the headfold, although its expression was not as consistent with the *Tfap2a* expression domain compared to *Gli3* (**Fig. S2C**). These data confirmed that Gli TFs were both temporally and spatially expressed in a manner consistent with a role in early NCC development.

### Early knockdown of GLI3 leads to reduced expression of neural crest cell specifiers

Given the spatiotemporal expression of *Gli3*, we next wanted to assess its role in developing NCCs. Knockdown experiments were first performed in chick embryos via unilateral electroporation with GLI3 morpholinos (GLI3-MO) at HH4, and subsequent analysis at ∼HH10. Electroporation with a control morpholino did not result in any changes in *PAX7* expression (**Fig. 3A**). Conversely, electroporation with a GLI3-MO resulted in reduction of *PAX7* expression (**Fig. 3B, white arrows)**. To test if GLI3 was similarly required for human NCC specification, we performed GLI3 knockdown in ES cell-derived human NCCs. Human ES cells (Day 0) were treated with either control Cy3- or *GLI3*-siRNA prior to NCC differentiation. Efficacy of *GLI3* knockdown was confirmed by qPCR (**Fig. S3A**). NCC differentiation was carried out and Day 5 NCCs were analyzed for expression of NCC markers including SOX10 and PAX7. Upon treatment with Cy3-siRNA, NCCs robustly expressed both SOX10 and PAX7. In contrast, siRNA knockdown with *GLI3*-siRNA led to a reduction in SOX10 and PAX7 protein expression (**Fig. 3C-D**). qPCR on *GLI3*-siRNA treated NCCs revealed reduced gene expression of *SOX10*, *PAX7, FOXD3* and *TFAP2A.* Together, these data confirmed that *GLI3* knockdown reduced expression of multiple NCC specifiers (**Fig. 3E**).

**Figure 3:**
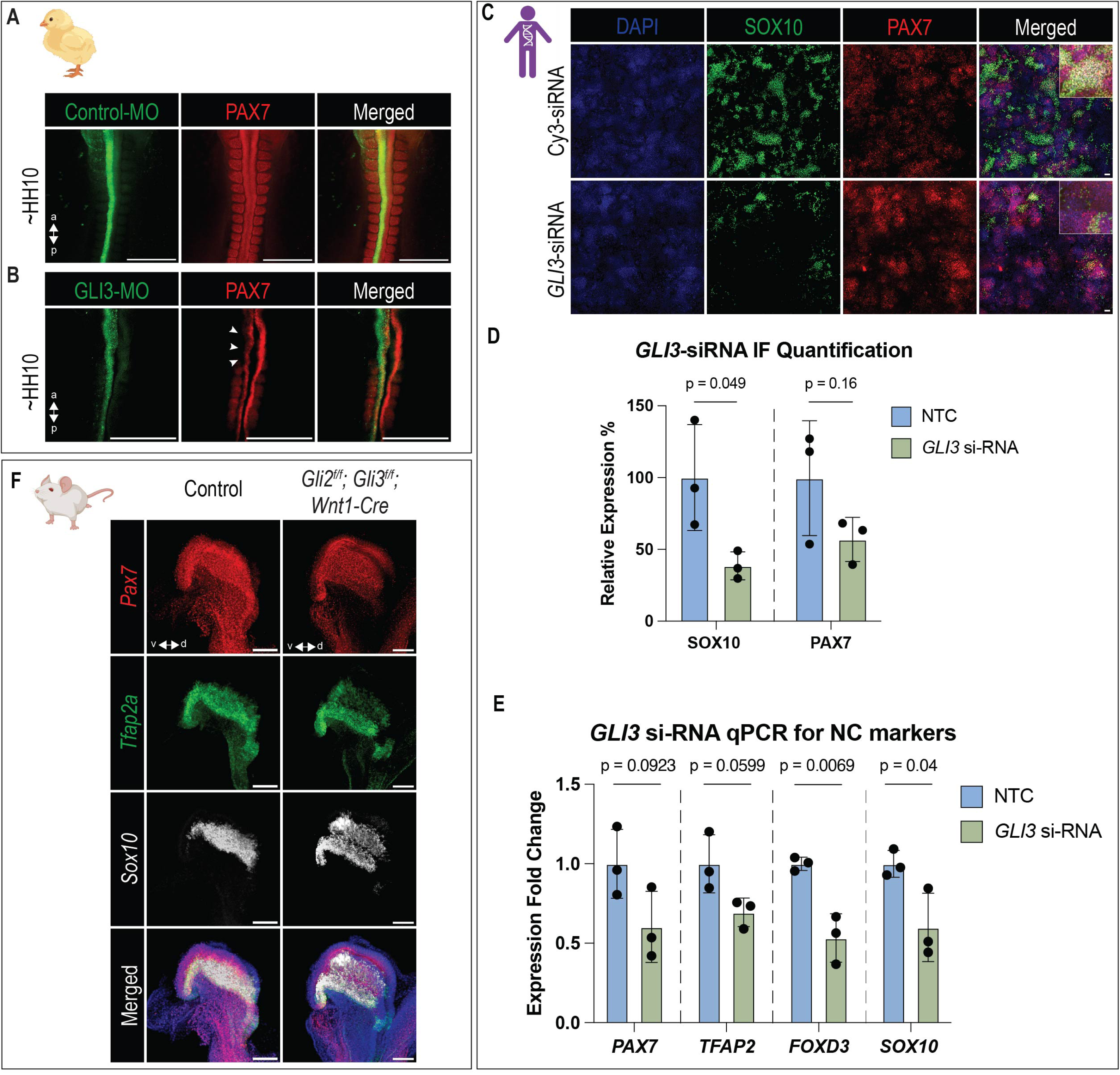
Early knockdown of GLI3 leads to reduced expression of neural crest cell specifiers. (A-B) ∼HH10 chick embryos injected with either (A) control or (B) GLI3 morpholinos stained for PAX7 protein expression. White arrows indicate reduced PAX7 expression. (C) Immunofluorescence of Day 5 hNCCs treated with either non-target control (NTC, Cy3-siRNA) or *GLI3*-siRNA showing expression of neural crest markers SOX10 and PAX7. Inset displays higher magnification. (D) Quantification of immunofluorescence from (C). (E) qPCR for neural crest specification genes in hNCCs treated with either non-target control (NTC, Cy3-siRNA) or *GLI3*-siRNA. (F) Whole-mount RNAscope of E8.25 control and *Gli2^f/f^; Gli3^f/f^; Wnt1-Cre* embryos showing expression of neural crest specification genes *Pax7, Tfap2a, and Sox10*. Animal model schematics were created using Biorender.com. Scale bars: 500μm (A, B), 50μm (C), 100μm (F).

To determine if GLI regulation of NCC specifiers was conserved in a murine model, the Cre-lox conditional knockout system was utilized. Since our work detected *Gli2* expression in NCCs and previous studies suggested redundancy of GLI TFs in the specification of NCCs (McDermott et al., 2005, Cerrizuela et al., 2018, Mo et al., 1997), we generated *Gli2^f/f^; Gli3^f/f^; Wnt1-Cre* embryos. Interestingly, *Gli2^f/f^; Gli3^f/f^; Wnt1-Cre* embryos did not present with a notable difference of *Pax7*, *Tfap2a*, or *Sox10* expression (**Fig. 3F, S3B**). While species-specific variation has been documented in NCC development (Barriga et al., 2015, Stuhlmiller and Garcia-Castro, 2012), this result was surprising and led to additional analysis of the temporal window of recombination of *Wnt1-Cre* (**Fig. S3C**). Characterization of *R26R-eYFP; Wnt1-Cre* embryos from E7.75-E8.25 supported the conclusion that recombination with the *Wnt1-Cre* driver occurred at ∼E8.0, a point at which NCC have already received induction and specification signals (Barriga et al., 2015). Thus, as other recent reports have suggested (Barriga et al., 2015, Moore Zajic et al., 2025, Debbache et al., 2018), we surmised that recombination with the *Wnt1-Cre* driver did not occur early enough to adequately test the hypothesis that loss of *Gli3* prevented initial expression of NCC specifier genes. Despite failure to observe loss of NCC specifiers, our previous work reported that conditional loss of Gli TFs using the *Wnt1-Cre* driver resulted in phenotypes consistent with impaired NCC development (Elliott et al., 2020).Thus, we next explored a second, later role for GLI3 in the differentiation of NCCs.

### GLI3 binds at promoter regions of genes associated with ectomesenchymal and non-ectomesenchymal cranial neural crest cell differentiation

In addition to robust expression in the dorsal neural tube, *Gli3* expression is maintained in facial prominences during cranial neural crest cell (CNCC) differentiation (Du et al., 2012, Chang et al., 2016) and has been implicated in development of the CNCC-derived craniofacial skeleton (Elliott et al., 2020). Despite these data, the mechanism by which GLI3 contributes to CNCC differentiation remains nebulous. To determine a role for GLI TFs in CNCC differentiation, E11.5 mandibular prominences from *3xFLAG-Gli3* embryos were harvested and Cleavage Under Targets and Release Using Nuclease (CUT&RUN) (Skene and Henikoff, 2017) was performed (**Fig. 4A**). A consensus peak set of GLI3 binding compared to IgG controls (n = 3,895) was defined based on presence in ≥2/3 biological replicates (**Fig. 4B, Fig. S4A**). *De novo* motif analysis identified the canonical GLI binding motif (GBM: GACCACC) (Peterson et al., 2012) as the top ranked motif, with the SP1 (zinc finger transcription factor) and NF-Y (CCAAT-binding factor) motifs as the top three (**Fig. 4C**). The GLI3 consensus peaks strongly enriched for promoter regions (>75%), with 12% being intergenic, 9% being intronic, and 2% being exonic, and the remainder found in other genomic regions (**Fig. 4D**). Gene Ontology (GO) term analysis enriched for both ectomesenchymal and non-ectomesenchymal GO terms (**Fig. 4E-F, Supplemental Table 1**). These results strongly suggested that GLI3 not only played a role in early NCC specification but also played a role later in CNCC differentiation.

**Figure 4:**
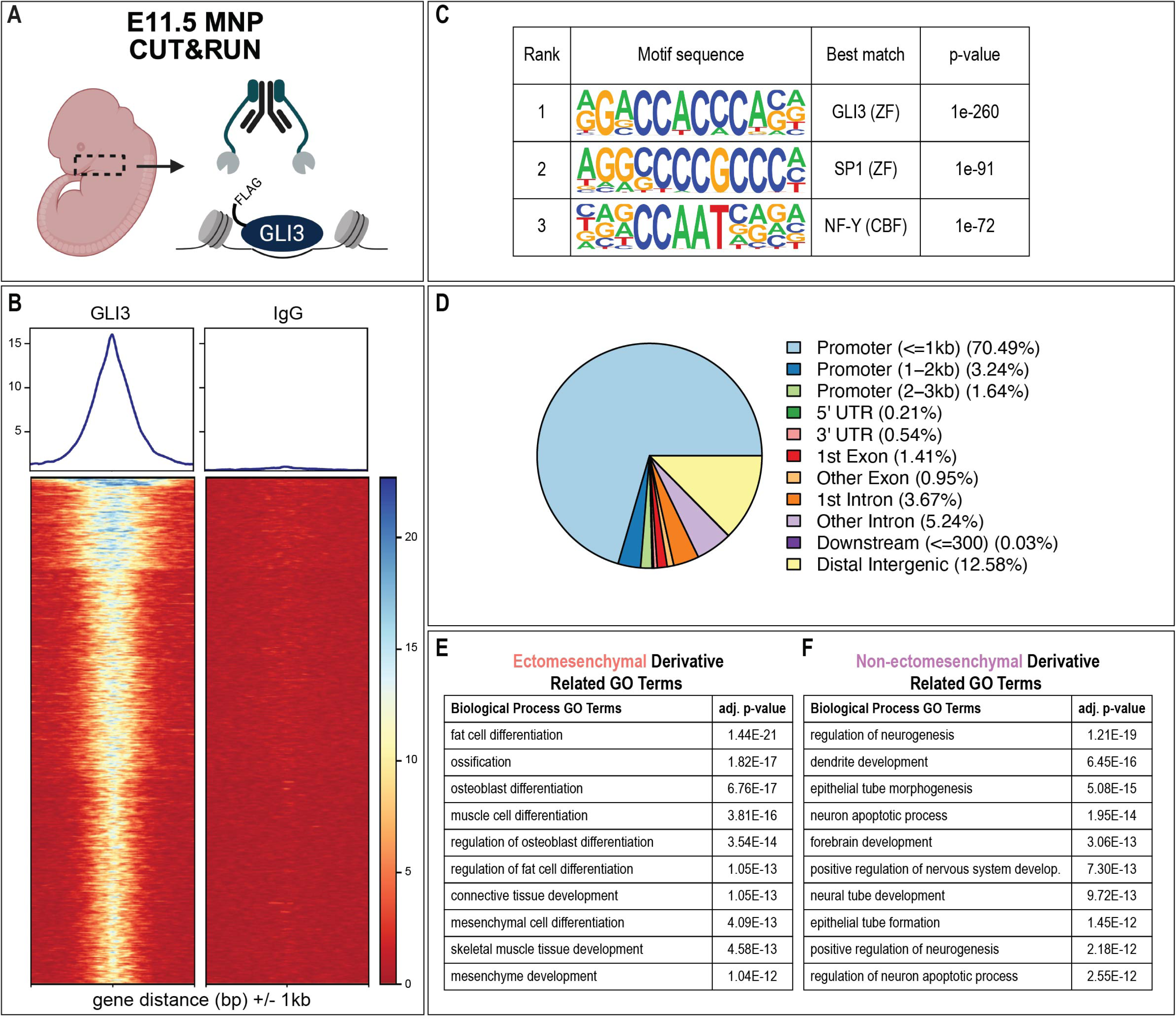
GLI3 binds at promoter regions of genes associated with ectomesenchymal and non-ectomesenchymal cranial neural crest differentiation. (A) Schematic of CUT&RUN experiment performed on mandibular prominences (MNPs) of E11.5 *3xFLAG-Gli3* embryos. (B) Heatmap of GLI3 bound regions compared to IgG control. (C) Top three motifs enriched in GLI3 bound regions determined by HOMER. (D) Distribution of GLI3 bound regions from CUT&RUN dataset at key genomic regions. (E-F) Relevant GO terms for (E) ectomesenchymal and (F) non-ectomesenchymal CNCC derivatives found in the top 50 GO terms ranked by adjusted p-value.

### GLI3 peaks are associated with active transcription of ectomesenchymal genes

While both GLI2 and GLI3 are bimodal transcription factors that activate and repress downstream targets, GLI2 is thought to function as the predominant activator, while GLI3 is thought to function as the predominant repressor (Park et al., 2000, Bai et al., 2004). To determine whether GLI3 binding was associated with target gene activation or repression in CNCCs, we used a publicly available ChIP-seq dataset for histone marks (H3K27ac, H3K27me3, and H3K4me1) from E11.5 mouse mandibular prominences (Visel et al., 2017). Alignment of GLI3 peaks with the histone mark datasets revealed that GLI3 binding peaks were highly associated with H3K27ac (**Fig. 5A**), a mark indicative of active transcription. K-means clustering was performed to yield 2 major clusters. Cluster 1 was strongly associated with H3K27ac, and motif enrichment analysis yielded similar motifs as those observed in GLI3 CUT&RUN analysis (**Fig. 5B and 4C**). GO term analysis of Cluster 1 showed terms associated with splicing, chromatin organization, stem cell population maintenance, and osteoblast differentiation (**Fig. 5C**). Cluster 2, contained the majority of GLI3 peaks, had lower associated with H3K27ac, yet utilized similar motifs as compared to Cluster 1 (**Fig. 5D**). GO term analysis of Cluster 2 revealed terms associated with Wnt signaling as well as terms associated with ossification and skeletal muscle development (**Fig. 5E**). Visualization of genes associated with GLI3 peaks revealed significant GLI3 binding at several genes involved in ectomesenchymal (**Fig. 5F-I**), but not non-ectomesenchymal differentiation (**Fig. 5J-M**). Taken together, these data suggest that GLI3 directly activated genes required for ectomesenchymal differentiation of CNCCs.

**Figure 5:**
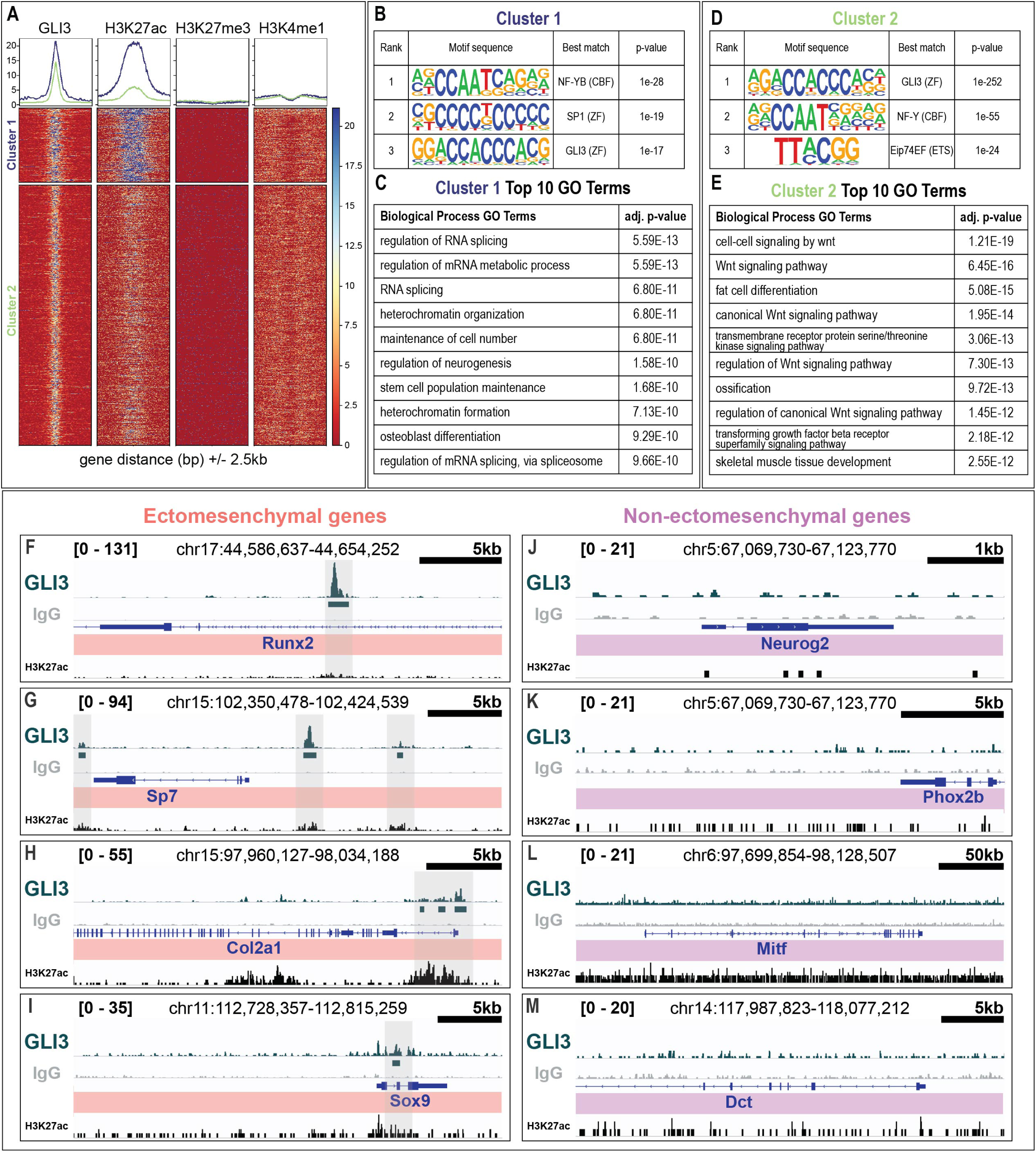
GLI3 peaks are associated with active transcription of ectomesenchymal genes. (A) Heatmap of GLI3 bound regions relative to publicly available H3K27ac, H3K27me3, and H3K4me1 ChIP-seq data. (B) Top three motifs enriched in Cluster 1 of GLI3 bound regions determined by HOMER. (C) Top 10 GO terms for Cluster 1. (D) Top three motifs enriched in Cluster 2 of GLI3 bound regions determined by HOMER. (E) Top 10 GO terms for Cluster 2. (F-M) IGV browser tracks of ectomesenchymal (F-I) and non-ectomesenchymal (J-M) genes. GLI3 peaks (green peak), GLI3 bound regions (green block lines), IgG control peaks (gray), loci (blue), and H3K27ac peaks (black).

### Loss of GLI TFs impairs differentiation of ectomesenchymal, but not non-ectomesenchymal cranial neural crest cell derivatives

To definitively test the role of GLI-mediated signaling during CNCCs differentiation, we examined development of CNCC derivatives in *Gli2^f/f^; Gli3^f/f^; Wnt1-Cre* embryos (Chang et al., 2016, Millington et al., 2017, Elliott et al., 2020). H&E staining of E13.5 control and *Gli2^f/f^; Gli3^f/f^; Wnt1-Cre* embryos revealed a loss or reduction of several structures that require CNCC contributions including the tongue, sub-mandibular glands, palatal shelves, teeth, and skeletal elements (**Fig. 6A-B**) (Millington et al., 2017, Elliott et al., 2020). Alizarin Red/Alcian Blue staining of E18.5 control and *Gli2^f/f^; Gli3^f/f^; Wnt1-Cre* heads revealed additional skeletal phenotypes including micrognathia and hypoplastic calvaria (**Fig. 6C-F, Fig. S5A-B)**. To confirm these phenotypes were due to the loss of GLI TFs in CNCCs, we repeated this experiment with two additional Cre lines that also target CNCCs later than *Wnt1-Cre*: *Pax3^Cre/+^* and *Sox10-Cre*. *Pax3* expression was first detected at ∼E8.5 in the dorsal neural tube during CNCC specification (Goulding et al., 1991) and has previously been utilized as an alternative Cre line for conditional gene knockout in pre-migratory NCCs (Engleka et al., 2005). *Sox10-Cre* recombines in NCCs at ∼E9.0 post delamination from the neural tube (Paratore et al., 2001, Hari et al., 2012) and is commonly used for conditional knockout in post-migratory NCCs (Matsuoka et al., 2005). Loss of *Gli* TFs in both *Pax3^Cre/+^* and *Sox10-Cre* embryos led to similar phenotypes as those observed in *Wnt1-Cre* embryos, such as loss or reduction of the tongue, sub-mandibular glands, palatal shelves and teeth (**Fig. S5C**). Alizarin Red/Alcian Blue staining of E18.5 *Gli2^f/f^; Gli3^f/f^; Pax3^Cre/+^* and *Gli2^f/f^; Gli3^f/f^; Sox10-Cre* embryos showed similar phenotypes as *Gli2^f/f^; Gli3^f/f^; Wnt1-Cre* such as micrognathia and a reduction in the frontal bone (**Fig. 6G-J, Fig S5D-E**). While the mandible was significantly shorter in all mutants compared to controls (**Fig. S5F**), the frontal bone width was only significantly decreased in *Gli2^f/f^; Gli3^f/f^; Pax3^Cre/+^* and *Gli2^f/f^; Gli3^f/f^; Wnt1-Cre* embryos (**Fig. S5G**), suggesting a temporal window for GLI function during calvaria morphogenesis.

**Figure 6.**
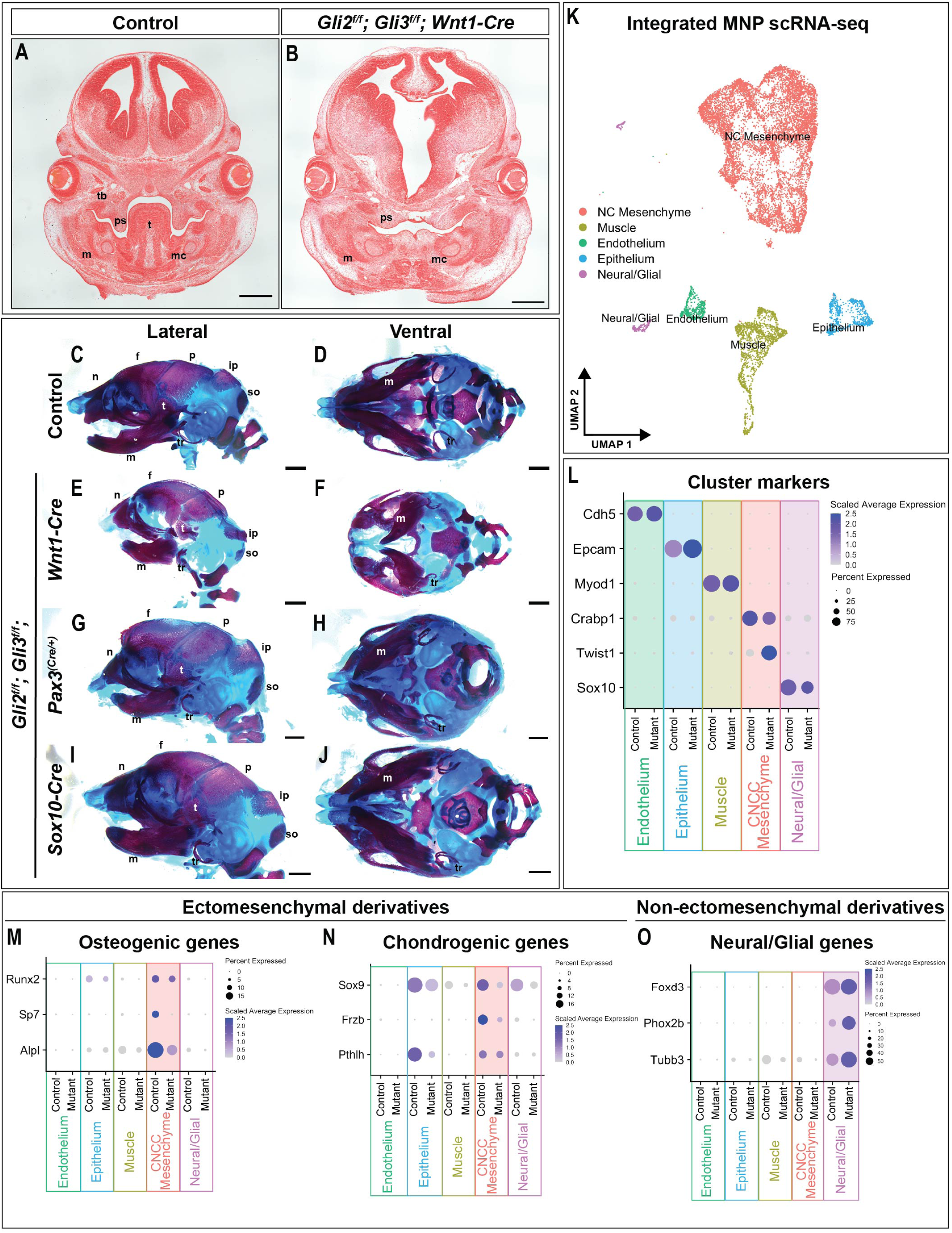
Loss of GLI TFs impairs ectomesenchymal, but not non-ectomesenchymal CNCC differentiation. (A-B) H&E staining of frontal sections from (A) *Cre*-negative control and (B) *Gli2^f/f^; Gli3^f/f^; Wnt1-Cre* E13.5 embryos. (C-J) Lateral and ventral views of Alcian Blue and Alizarin Red stained E18.5 (C-D) control, (E-F) *Gli2^f/f^; Gli3^f/f^; Wnt1-Cre*, (G-H) *Gli2^f/f^; Gli3^f/f^; Pax3^(Cre/+)^*, and (I-J) *Gli2^f/f^; Gli3^f/f^; Sox10-Cre* embryonic skulls. (K) UMAPs of integrated scRNA-seq data from control and *Gli2^f/f^; Gli3^f/f^; Wnt1-Cre* mandibular prominences (MNPs). (L) DotPlot of expression of genes used for clustering the integrated scRNA data of control and *Gli2^f/f^; Gli3^f/f^; Wnt1-Cre* mutant MNPs. (M-O) DotPlots of (M) osteogenic and (N) chondrogenic genes in control and *Gli2^f/f^; Gli3^f/f^; Wnt1-Cre* mutant MNPs. (O) DotPlot of neural/glial genes in control and *Gli2^f/f^; Gli3^f/f^; Wnt1-Cre* mutant MNPs. (f) frontal bone; (ip) interparietal bone; (m) mandible; (mc) Meckel’s cartilage; (n) nasal bone; (t) tongue; (tb) tooth bud; (p) parietal bone; (ps) palatal shelves; (so) supraoccipital bone; (tr) tympanic ring. Scale bars: 500µm (A, B), 1mm (C-J).

Multipotent CNCCs undergo various transcriptional changes to initiate differentiation into ectomesenchymal and non-ectomesenchymal derivatives (Le Lievre and Le Douarin, 1975). To determine how loss of GLI TFs affected the CNCC transcriptome, scRNA-seq was performed on E13.5 control and *Gli2^f/f^; Gli3^f/f^; Wnt1-Cre* mandibular prominences (MNPs). Integration of control and mutant mandibular prominences yielded 19 distinct clusters, grouped into five larger clusters by expression of cell type markers: Endothelium (*Cdh5*), Epithelium (*Epcam*), Muscle (*Myod1*), CNCC Mesenchyme (*Crabp1*), and Neural/Glial (*Sox10*) (**Fig. 6K, L; Fig. S5H, I**). Whereas minimal gene expression changes were observed between the control and mutant in markers used for clustering endothelium, epithelium and muscle, significant changes were observed in mesenchymal and neural clusters (**Fig. 6L**). Expression of *Twist1*, a TF that mediates cell fate choices through functional interactions with other proteins (Fan et al., 2021) was increased in *Gli2^f/f^; Gli3^f/f^; Wnt1-Cre* embryos, suggesting that loss of GLI TFs prevented the initiation of gene regulatory networks associated with differentiation. To further test if loss of GLI TFs impaired differentiation, downstream transcriptional regulators of ectomesenchymal and non-ectomesenchymal cell types were plotted. Genes associated with osteogenic differentiation such as *Runx2*, *Sp7, and Alpl* were decreased in mutant embryos (**Fig. 6M**). Similarly, genes associated with chondrogenic differentiation such as *Sox9* and *Fzrb* were also decreased in mutant embryos (**Fig. 6N**). In contrast, neural/glial markers such as *Foxd3*, *Phox2b*, and *Tubb3* were upregulated in *Gli2^f/f^; Gli3^f/f^; Wnt1-Cre* embryos (**Fig. 6O**), suggesting that loss of GLI TFs in CNCCs affected differentiation into ectomesenchymal but not non-ectomesenchymal derivatives. Together, the reduction of ectomesenchymal structures along with decreased expression of ectomesenchymal transcriptional regulators suggested that GLI TFs played a role in the ability of CNCC to differentiate into skeletal derivatives.

## Discussion

This study characterized previously unappreciated roles for GLI3 in NCC development. Examining three different species (i.e., chick, mouse, and human) we demonstrated that *GLI3* was co-expressed with several well established NCC specifier genes during NCC induction/specification and knocking down *GLI3* expression prior to NCC specification resulted in the downregulation of NCC specifiers (**Fig. 7A**). Furthermore, conditional knockout of GLI TFs after NCCs were specified revealed a second role for GLI during NCC differentiation, in which GLI3 activity was required for ectomesenchymal (e.g., bone/cartilage) differentiation (**Fig. 7B**). Interestingly, differentiation of other NCC derivatives (e.g., neurons) was not affected by the loss of *Gli3*. Our work builds off, and expands upon, previous studies that suggest a role for Shh/Gli in the induction, migration, and differentiation of NCCs in *Xenopus* and zebrafish embryos (Cerrizuela et al., 2018, Wada et al., 2005).

**Figure 7:**
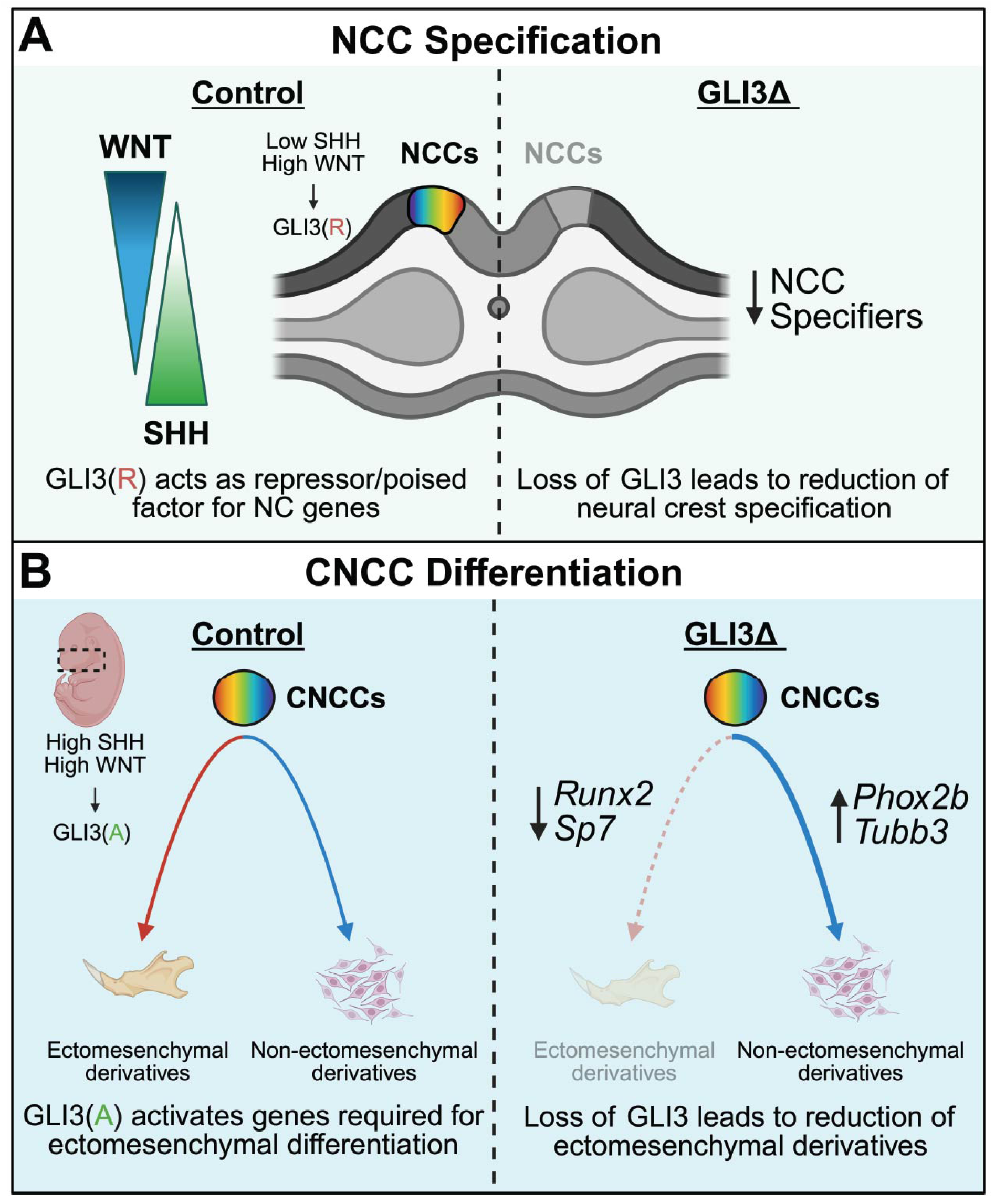
GLI3 functions during neural crest specification and differentiation. (A) Schematic of a cross-section through an embryo during NCC specification. GLI3 is induced by SHH or WNT signaling and functions to regulate expression of NCC specifiers. (B) Cranial neural crest cells differentiate into both ectomesenchymal and non-ectomesenchymal derivatives. Loss of GLI3 negatively affects ectomesenchymal, but not non-ectomesenchymal differentiation. Figure created using Biorender.com.

SHH pathway signal transduction is mediated by GLI transcription factors: GLI1, GLI2, and GLI3. GLI1, unlike GLI2 and GLI3, does not contain a repressor domain (Dai et al., 1999). GLI1 has also been shown to be dispensable in mice (Park et al., 2000), suggesting it may not be required for SHH signaling, but rather as an potentiator of the pathway (Sasaki et al., 1999, Dai et al., 1999). Conversely, GLI2 and GLI3 are bimodal and can act as either full-length activators (GLIA) or truncated repressors (GLIR) to mediate downstream targets (Sasaki et al., 1999). It is commonly accepted that GLI2 exists predominantly as a activator and GLI3 exists predominantly as a repressor due to protein isoform stability; however, there have been examples where GLI2 repressor activity and GLI3 activator activity have been identified (Bai et al., 2004, Wang et al., 2007, Bowers et al., 2012, Pan et al., 2009, Elliott et al., 2020). While our data do not explicitly distinguish between GLI3A and GLI3R either in DNA-binding or in transcriptional activity, we hypothesize that GLI3R may be the predominant isoform during early NCC specification, whereas GLI3A may be the predominant isoform during ectomesenchymal differentiation (**Fig. 7**).

*Gli3* expression is localized to the dorsal aspect of the neural tube during neurulation, prior to NCC specification (Persson et al., 2002). The expression of *Gli3* is complimentary to expression of *Shh* and *Gli2* in the ventral aspect of the neural tube. This complimentary expression establishes a HH gradient in which robust HH activity ventrally contributes to motor neuron subtype specification. Thus, these data suggest that it is the action of the GLI3R that establishes this gradient (Persson et al., 2002). While our data showing robust amounts of GLI3R in human NCCs supports this supposition, we have not explicitly determined that the repressor isoform is binding and repressing targets that must be downregulated for NCC induction and specification within the spatiotemporal window of NCC induction. Distinguishing between GLI3A and GLI3R isoforms utilizing a dual-tagged lines is a focus of our ongoing and future work.

While we hypothesize GLI3R is the predominant isoform during NCC specification, data herein supported a role for GLI3A during differentiation of cranial NCCs into ectomesenchymal derivatives. These data are in alignment with our previous findings which showed GLI3A activity was essential for mandibular development, via engaging with co-factors and utilizing divergent, low affinity motifs (Elliott et al., 2020). In contrast to the neural tube, expression of *Gli3* in the mandibular prominence is in the vicinity of *Shh* expression, and we observed marks of active transcription at targets required for skeletal differentiation (e.g., *Runx2, Sox9*). Despite these data, it remains possible that this transcription was driven by de-repression rather than activation, as has been proposed in the limb (Lex et al., 2022).

Regardless of isoform predominance, GLIs are well established as transcriptional drivers of the HH pathway (Sasaki et al., 1999). While this is indeed the case, several lines of evidence have also been presented suggesting that *Gli3*, in particular, is a target of the WNT pathway (Hasenpusch-Theil et al., 2012, Borycki et al., 2000, Alvarez-Medina et al., 2008, Mullor et al., 2001, Ulloa et al., 2007). With robust *Wnt* expression in the dorsal neural tube, it is likely that WNT signaling induces *Gli3* expression, which contributes to the establishment of the dorsal/ventral gradient of HH activity. It remains to be determined if induction by WNT, as opposed to induction by SHH, contributes to the eventual predominance of repressor versus activator isoforms. Furthermore, many studies have explored the convergence of these signaling pathways. Thus, it is possible that induction and function of GLI is affected by inputs from both pathways.

While NCCs historically have been referred to as multipotent (Bronner and LeDouarin, 2012), cranial NCCs have recently been given an elevated moniker of being “pleistopotent”, to describe their ability to differentiate into cells from more than one germ layer (Perera and Kerosuo, 2021). Pleistopotency is a distinct feature of cranial NCCs, as they are induced and specified as an ectodermal population yet give rise to cell-types traditionally attributed to the mesoderm, such as bone and cartilage. The ability of cranial NCCs to differentiate into ectomesenchymal derivatives is perhaps one of the most intriguing qualities of this cell type, and there have been several hypotheses proposed to explain how this potency is established (Buitrago-Delgado et al., 2015, Zalc et al., 2021, Perera and Kerosuo, 2021, Moore Zajic et al., 2025, Pajanoja et al., 2023). Previous *in vitro* studies demonstrated that exposure to recombinant SHH protein enhanced cranial NCC potency by increasing the proportion of cranial NCCs that differentiated into both non-ectomesenchymal and ectomesenchymal derivatives and decreased the proportion of neural-restricted precursors (Calloni et al., 2007). Furthermore, exposure to recombinant SHH protein promoted the development of cartilage from trunk neural crest cells (Calloni et al., 2007), a population of cells that does not normally give rise to ectomesenchymal derivatives. Numerous examples exist in the literature linking SHH signaling to potency and stemness in other cell types (Ishikawa et al., 2017, Ishikawa Y et al., 2021, Rowton et al., 2022, Yang et al., 2019). Despite these examples, the mechanism behind how SHH signaling led to an increased potency in CNCCs remains unclear. While our experiments did not directly test this hypothesis it remains possible exposure to SHH prior to induction and specification, and subsequent expression of GLI TFs, could contribute to CNCC potency and the ability to give rise to ectomesenchymal derivatives. Exploring the role of GLI TFs in CNCC potency is an ongoing line of research in our laboratories.

## Materials and Methods

### Analysis of publicly available single-cell RNA-sequencing and bulk RNA-sequencing datasets

For the analysis of RNA-sequencing dataset of sorted chick NC (Hovland et al., 2022), genes were plotted using the authors’ public R Shiny app, accessible at http://ash274.shinyapps.io/RNA-Seq_App/). Plots were generated with genes of interest and colors were modified using Adobe Illustrator.

For the analysis of single-cell RNA sequencing of early chick embryos (Perera and Kerosuo, 2021), the counts matrices were downloaded from the Gene Expression Omnibus (accession no. GSE221188). Seurat v4.3.0 (Hao et al., 2021) was used for downstream analysis. Cells were processed as stated in the authors’ publicly available scripts (https://github.com/KerosuoLab/Pajanoja_2023). A DotPlot was generated for the genes from the publication to determine similarity between the original and reanalyzed datasets. DotPlots of genes of interest were then generated.

For the analysis of single-cell RNA-sequencing dataset of sorted CNCCs (Zalc et al., 2021), the counts matrix was downloaded from the Gene Expression Omnibus (accession no. GSE162035). Cells for downstream analysis were subset by the following criteria: more than 500 features, less than 25,000 features, more than 50,000 counts, less than 200,000 counts, and less than 5% mitochondrial gene expression. Cells were then taken through the standard Seurat processing which included *NormalizeData, FindVariableFeatures, and ScaleData*. Cells were clustered by *RunPCA*, *FindNeighbors*, *FindClusters*, *RunUMAP*, and *RunTSNE.* A DotPlot was generated for the genes from the publication to determine similarity between the original and reanalyzed datasets. Clusters were then renamed and reordered to best match the original published dataset. DotPlots of genes of interest were then generated.

### Animal husbandry and embryo collection

For mouse husbandry, embryo collection, and tissue preparation both male and female mice were used. All mouse protocols used were approved by the Institutional Animal Care and Use Committee (IACUC) and maintained by the Division of Veterinary Services at Cincinnati Children’s Hospital Medical Center.

Timed matings were performed with noon of the day a vaginal plug was discovered designated as E0.5 Embryos were harvested between E7.5-E18.5, collected and harvested in DEPC PBS. Embryos were fixed in 4% paraformaldehyde from 1 hour at room temperature to overnight at 4°C based on stage. Embryos for whole-mount RNAscope were dehydrated through a methanol series and kept in -80°C. For paraffin embedding, embryos were dehydrated through an ethanol series, washed in xylene, and embedded in paraffin. All mouse embryo experiments were performed with a minimum of three biological replicates unless indicated otherwise.

For chicken husbandry, embryo collection, and tissue preparation, fertilized White Leghorn chicken eggs were obtained from Sunstate Ranch (Sylmar, CA) and incubated at 37 °C under humidified conditions until they reached the desired developmental stage, as defined by the Hamburger and Hamilton (HH) staging system (Hamburger and Hamilton, 1951). Embryos at the appropriate HH stages were isolated and fixed in 4% paraformaldehyde (PFA) in phosphate-buffered saline (PBS) for 10-20 minutes, depending on the developmental stage. Fixed embryos were subsequently processed for *in situ* hybridization and immunofluorescence to localize *GLI3* and *PAX7* mRNA and protein expression, respectively. All chick embryo experiments were performed with a minimum of three biological replicates unless indicated otherwise.

### Chick embryo in situ, immunofluorescence, and morpholino experiments

Whole mount chicken embryos were used for *in situ* hybridization. An anti-sense mRNA probe targeting *GLI3* mRNA was created using T7 Reverse Transcription from chicken cDNA template generated via PCR. Sequences for forward and reverse primers are listed in Supplemental Table 2. Color reaction generated via NBT/BCIP substrate reaction with DIG labeled antibody. Expression was assessed with minimum 5 embryos per stage with representative images shown.

For IF, embryos were blocked for 2 hours with 10% Donkey Serum in PBS Tween. GLI3 antibody (R&D Systems, AF3690) and PAX7 antibody (DSHB, AB_528428) incubated overnight at 4 °C. 1:2000 dilutions of secondary antibodies for GLI3 (Alexafluor Donkey anti-Goat 488) and for PAX7 (Alexafluor Donkey anti-Mouse 647) were used and embryos were incubated overnight at 4°C. Expression was assessed with a minimum of 5 embryos per stage and imaged on an Inverted Nikon Elements.

For chick embryo morpholino injection fertilized eggs were incubated at 37°C in 60% relative humidity until they reached HH4. Embryos at HH4 were isolated, and a GLI3 morpholino 5’-GCCTCCATGATGTCTTCTCATTACT-3’-FITC and Control morpholino 5’-CCTCTTACCTCAgTTACAATTTATA-3’-lissamine (Gene Tools, LLC) was injected at a final concentration of 1 mM. Morpholinos were prepared in a 30% sucrose solution mixed at a 1:1 ratio. Electroporation was performed using a B.T.C. electroporation system with five pulses of 50 ms at 5.1 V with an interval of 100ms between pulses (Williams and Sauka-Spengler, 2021). Following electroporation, embryos were cultured *ex ovo* using the Easy Culture method until the desired developmental stage (Chapman et al., 2001). Electroporated embryos were immunostained for PAX7 (DSHB, AB_528428).

### Human embryonic stem cell (hESC) and neural crest cell (hNCC) experiments

H1 embryonic stem cells (WA01 WiCell Institute) were grown and maintained on Matrigel coated plastic plates using mTesr1 (STEMCELL Technologies, 85850). Cultures were passaged with Versene (Lonza) every 4-5 days with Passage 26-35 being used for neural crest differentiation. For neural crest induction, hESC cells were dissociated with Accutase (STEMCELL Technologies) at 70-80% confluency. Single cells were resuspended in Induction media supplemented with 2.75µm CHIR 99021 and 10µm ROCK Inhibitor (STEMCELL Technologies, Y-27632). Induction media contains DMEM/F12 with 2% B27, 1x Glutamax (Gibco, 35050061) and 0.5% BSA. Cells were plated at 20k/cm^2^ in Induction media supplemented with CHIR and ROCK inhibitor for the first 2 days. Induction media without CHIR and ROCK inhibitor was used for the remaining 3 days (Leung et al., 2016).

RNA-sequencing data was downloaded from the Gene Expression Omnibus (accession no. GSE125145). Briefly, sequencing reads were trimmed for low quality bases using sickle v1 (https://github.com/najoshi/sickle). Trimmed reads were mapped to the hg38 reference genome using STAR version 2.7.9 (Dobin et al., 2013). Count matrix of reads was obtained using Subread 2.0.3 (Liao et al., 2014) and differential expression analysis was completed using DESeq2 (Love et al., 2014). Heatmaps were generated using the ComplexHeatmap (Gu et al., 2016) using scaled, VST-normalized counts.

For western blot analysis of human neural crest cells, human embryonic stem cells and induced neural crest cells at 80–90% confluency were detached from culture wells using Accutase for 3 minutes. Cells were then resuspended in DMEM and centrifuged to obtain cell pellets. The pellets were subsequently washed with PBS and stored at -80 °C prior to protein isolation. Western blots were performed as described in (Millington et al., 2017). Briefly, samples were sonicated in RIPA buffer containing protease and phosphatase inhibitors. Protein concentration was measured by BCA protein assay. 10μg of protein was loaded to a NuPAGE^TM^ Tris-Acetate Mini Protein Gel (ThermoFisher, EA03785BOX). Antibodies used: anti-GLI3 (1:1000; R&D Systems, AF3690), anti-Vinculin (1:2000; Santa Cruz Biotechnology, sc-73614), IRDye^®^ 800CW Donkey anti-Rabbit IgG (1:2000; LICOR, 926-32213), and IRDye^®^ 680RD Donkey anti-Mouse IgG (1:2000; LICOR, 925-68072). Images were taken by LICOR Odyssey^®^ DLx.

siRNA was performed as described in (Prasad et al., 2020) using si-RNA for GLI3 (Dharmacon L-011043-00-0005) and Silencer Cy3 labeled Negative Control (Thermofisher AM4621). Assessment of knockdown efficiency was done at Day2 with qRT-PCR. si-RNA was reverse transfected with Lipofectamine RNAiMAX (Thermofisher 13778030). si-RNA and RNAiMAX was mixed with OptiMEM and incubated for 10-20 minutes before plating on Matrigel coated 96 well plate. Media was changed daily with cells collected for RNA at Day 2 and Day 5 and immunofluorescence at Day 5.

Immunofluorescence was performed on 96-well plate as described in (Leung 2016). Primary antibody for PAX7 (DSHB, AB_528428) and SOX10 (Santa Cruz Biotechnology Inc., sc-271163). Secondary antibodies were at 1:2,000 including Goat anti-mouse IgM-488 (Thermo-Fisher A-21042), Goat anti-mouse IgG1-568 (Thermo-Fisher A-21124), Donkey anti-goat 647 (Thermo-Fisher A-21447), Donkey anti-mouse 488 (Thermo-Fisher A-21202). Images were taken with Nikon Eclipse Ti microscope and analyzed using Nikon Elements Software. Images processed in FIJI.

For gene expression analysis of siRNA treated human NCC, RNA was collected in TRIzol and purified with Directzol RNA Microprep kit (Zymo Research, R2061). RNA was reverse transcribed with PrimeScript Reverse Transcriptase kit (Takara, 2680A).

Quantitative PCR (qPCR) was done using TB Green Premix Ex Taq II Tli RNase H Plus (Takara, RR820A) on Applied Biosystems QuantStudio 6 Real-Time PCR System.

Primers were used at 300nM (Supplementary Table 2). ΔΔCT method was used to analyze expression relative to Cy3 Non-Targeting Control. Three to five biological replicates were used for statistical analysis with GAPDH used to normalize and reference gene.

### Whole-mount RNAscope in situ hybridization of mouse embryos

Whole mouse embryos were stained with ACD’s RNAscope Multiplex Fluorescent Kit v2.0 according to their “RNAscope Assay on whole-mount zebrafish embryos” technical note with modifications for E7.5-8.5 mouse embryos. Briefly, embryos were rehydrated through a methanol series, target retrieval and protease treatment were performed, and probes were hybridized overnight. The following day, embryos were taken through the AMP series, HRP series paired with fluorophores, then were placed in DAPI overnight. The next day, DAPI was washed out, and embryos were dehydrated through a methanol series before being mounted in RIMS (Yang et al., 2014) for clearing. Embryos were imaged with a Nikon A1 inverted confocal microscope system. Z-stacks of images were projected into 3D projections using Fiji (v.2.16.0) (Schindelin et al., 2012). A minimum of three independent biological replicates were used for each staining.

### Cleavage under targets and release using nuclease (CUT&RUN)

CUT&RUN was performed on 3 biological replicates according to the EpiCypher CUTANA CUT&RUN Protocol (v1.7) with modifications (Lex et al., 2022). Reads and data processing details for each sample can be found in NCBI’s Gene Expression Omnibus (GEO) under accession number GSE301704. Briefly, E11.5 mandibles were dissected in ice-cold PBS and dissociated in 100 μg/mL Liberase at 37°C. Samples were assessed for cell quality using Trypan Blue. Samples were then bound to activated ConA beads before being incubated overnight at 4°C with either 1:100 anti-FLAG M2 antibody (Sigma Aldrich, F1804) or IgG (R&D Systems, MAB002R). Samples were then incubated with secondary antibody and washed in digitonin wash buffer before incubation with CUTANA pAG-MNAse. DNA fragments were then purified before library preparation (Skene et al., 2018). Library preparation was conducted using NEBNext Ultra II DNA Library Prep Kit for Illumina (NEB, #E7645S). Samples were sequenced on an Illumina NextSeq500 instrument using EF-75 bp to a depth of 3-5 million reads.

Quality-controlled FASTQ files for each dataset were cleaned using Trim Galore version 0.6.6 (https://github.com/FelixKrueger/TrimGalore) to remove remaining adapter sequences. Reads were aligned to the mm10 genome using BowTie2 version 2.4.2 (Langmead and Salzberg, 2012) and duplicates were removed using SAMtools version 1.18.0 (Danecek et al., 2021). Reads overlapping blacklisted regions (Amemiya et al., 2019) were removed using BEDtools version 2.30.0 (Quinlan and Hall, 2010). Peaks were called against IgG control antibody samples using MACS 3.0.2 (Zhang et al., 2008). A consensus peak set for the anti-Gli3 datasets was generated using BEDtools, requiring peak presence in two of three biological replicate peak sets.

Single replicates of publicly available histone modification ChIP-seq datasets (H3K4me3, H3K27ac, H3K27me3) for E11.5 dissected mandibular prominences were obtained from FaceBase (https://www.facebase.org) (Samuels et al., 2020, Visel et al., 2017) and processed as above.

HOMER de novo motif analysis used version 4.9 (Heinz et al., 2010) to scan for 6, 12 and 20bp motifs in a 200bp region around each peak center. Top enriched motifs were compared to the HOMER known motif database and selected based on similarity. CUT&RUN peaks were annotated to the nearest transcription start site (TSS) using the ChIPseeker package in R (Wang et al., 2022, Yu et al., 2015). Gene Ontology (GO) term enrichment analysis for biological processes (BP) was conducted using clusterProfiler (Xu et al., 2024). Terms for display were selected for biological relevance or rank as indicated. bigWig files and heatmaps of read enrichment around CUT&RUN peaks were generated using deepTools2 version 3.5.6 (Ramirez et al., 2016). Gli3 CUT&RUN and histone modification ChIP-seq datasets were clustered around consensus CUT&RUN peaks using the K-means algorithm in deepTools, selecting K-means = 2 as the best separation of low- and high-level histone modification. CUT&RUN peak annotation was completed as above.

Processed tracks were visualized by loading them into IGV (v.2.14.0) (Robinson et al., 2011, Thorvaldsdottir et al., 2013) using the mm10 (GRCm38) mouse genome and edited with Adobe Illustrator.

### Mouse single-cell RNA sequencing

E13.5 mandibles from control CD1 or *Gli2^f/f^; Gli3^f/f^; Wnt1-Cre* mouse embryos were dissected in ice-cold DEPC PBS and minced to a fine paste. 12.5 mg of tissue was placed in a sterile 1.5 mL tube containing 0.5 mL protease solution containing 125 U/mL DNase and 5 mg/mL Bacillus Licheniformis. The samples were incubated at 4°C for a total of 10 minutes, with trituration using a wide boar pipette tip every minute after the first two. Protease was inactivated using ice-cold PBS containing 0.02% BSA and filtered using 30 mM filter. The cells were pelleted by centrifugation at 200g for 4 minutes and resuspended in 0.02% BSA in PBS. Cell number and viability were assessed using a hemocytometer and trypan blue staining. 9,600 cells were loaded onto a well on a 10X Chromium Single-Cell instrument (10X Genomics) to target sequencing of 6,000 cells. Barcoding, cDNA amplification, and library construction were performed using the Chromium Single-Cell 3’ Library and Gel Bead Kit v3. Post cDNA amplification and cleanup was performed using SPRI select reagent (Beckman Coulter, Cat# B23318). Post cDNA amplification and post library construction quality control was performed using the Agilent Bioanalyzer High Sensitivity kit (Agilent 5067–4626). Libraries were sequenced using a NovaSeq 6000 and the S2 flow cell. Sequencing parameters used were: Read 1, 28 cycles; Index i7, eight cycles; Read 2, 91 cycles, producing about 300 million reads. The sequencing output data was processing using CellRanger (http://cloud.10xgenomics.com) to obtain a gene-cell data matrix.

In each dataset, cells with less than 200 features, more than 6000 features, and more than 10% mitochondrial gene expression were removed. The remaining cells were then processed through the standard Seurat processing which included *NormalizeData, FindVariableFeatures, CellCycleScoring, and ScaleData*. Cells were clustered by *RunPCA*, *FindNeighbors*, *FindClusters*, and *RunUMAP*, and visualized by Uniform Manifold Approximation and Projection (UMAP) plots. The control and mutant datasets were then integrated using *FindIntegrationAnchors and IntegrateData*. The integrated dataset was then scaled and clustered as before to generate the integrated R object.

Clusters were annotated using genes determined by *FindConservedMarkers* for each cluster as well as known gene markers. Clusters were then pseudobulked into larger cell-type specific clusters (NC Mesenchyme, Epithelium, Endothelium, Muscle, Neural/Glial, and Hematopoietic). Hematopoietic clusters identified by *Hba-a1* and *Ptprc* were excluded from downstream analysis. Gene expression analyses were conducted by splitting the integrated dataset by its condition (Control, Mutant), then generating *DotPlots* for relevant genes (e.g., ectomesenchymal genes, non-ectomesenchymal genes.)

### Whole-mount skeletal staining and imaging

E18.5 murine embryos were harvested and blanched to facilitate removal of skin and as much soft tissue as possible. Embryos were then immersed in 100% ethanol for 48 hours, followed by acetone for 48 hours. Cartilage was then stained by 0.015% Alcian Blue solution (in 20% glacial acetic acid and 80% 200 proof ethanol) for 24 hours, followed by a 100% ethanol wash for 24 hours. Embryos were then immersed in 1% KOH, followed by 0.005% Alizarin Red (in 1% KOH) for 24 hours to stain for bone. Samples were then taken through a graded 1% KOH/glycerol solution series and maintained in 50% glycerol for imaging.

## Acknowledgements

This research was made possible, in part, using the Cincinnati Children’s Bio-Imaging and Analysis Facility [RRID:SCR_022628], the Cincinnati Children’s Genomics Sequencing Facility [RRID:SCR_022630], the Cincinnati Children’s Bioinformatics Core Facility [RRID:SCR_022622], as well as the Cincinnati Children’s Single Cell Genomics Facility [RRID:SCR_022653]. We specifically acknowledge the assistances of Sarah McLeod, Dr. Matthew Kofron, Kelly Bailie, and Praneet Chaturvedi. We would also like to thank the Cincinnati Children’s Division of Veterinary Services for their considerate care of the mouse colony. Finally, we thank members of the Brugmann Lab, the Gebelein Lab, and García-Castro Lab for their valuable discussions, comments, and feedback.

## Data and resource availability

All relevant data and details of resources can be found within the article and its supplementary information.

## Supplemental Figure Legends

**Supplemental Figure S1:**
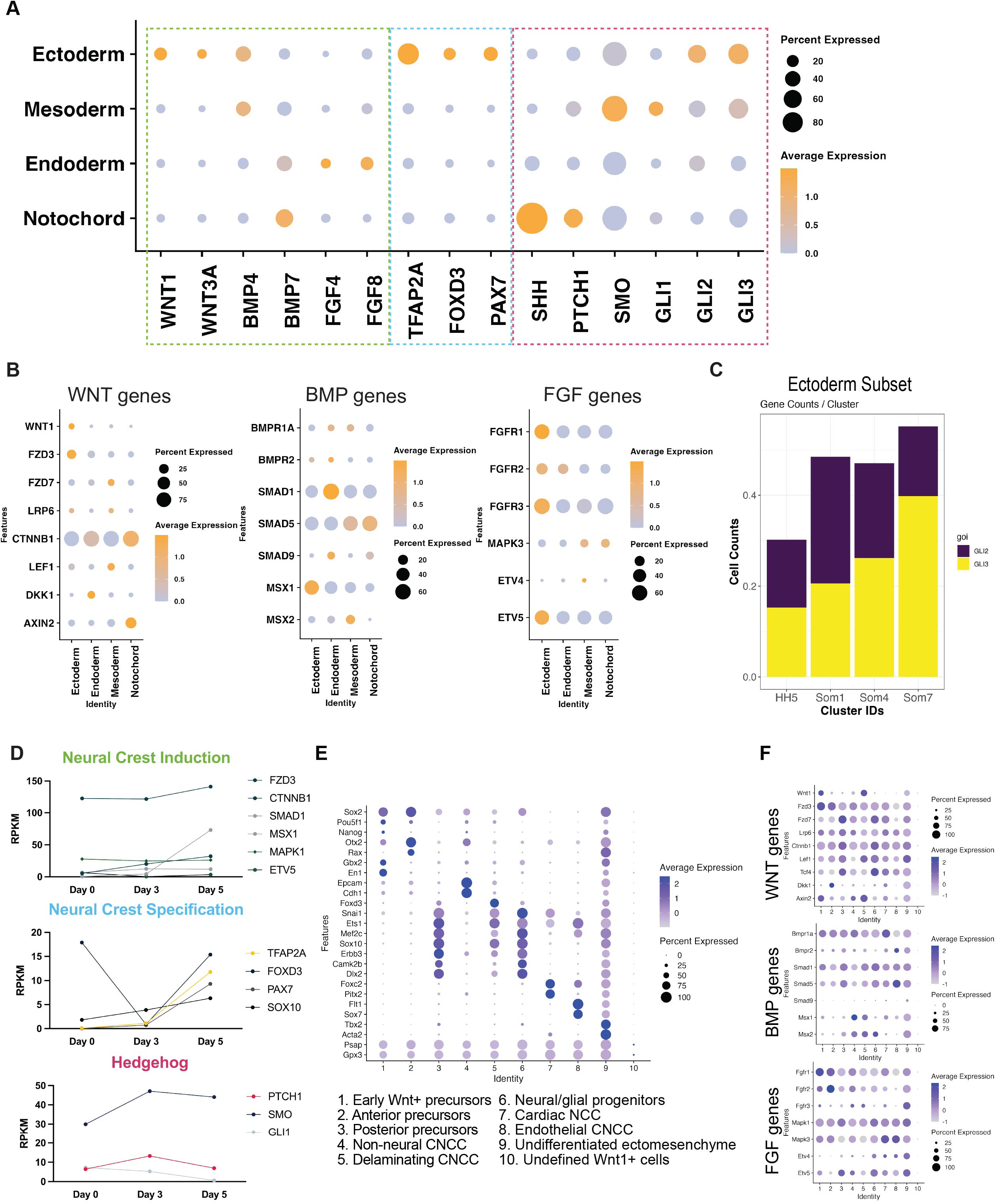
HH family members, including GLI3, are expressed in developing neural crest cells. (A) DotPlot of a scRNA-seq dataset from early chick embryos clustered by germ layer, reanalyzed to show expression of neural crest induction genes, neural crest specification genes, and SHH pathway genes. (B) DotPlots of additional WNT, BMP, and FGF family members from scRNA-seq dataset shown in (A). (C) Gene counts of GLI TFs from ectodermal cluster through chick embryonic development. (D) Reads per kilobase million (RPKM) of genes from heatmap in Figure 1B. (E) Full replication of original scRNA-seq dataset used in Figure 1D. (F) DotPlots of additional WNT, BMP, and FGF family members from scRNA-seq dataset shown in Figure 1D.

**Supplemental Figure S2:**
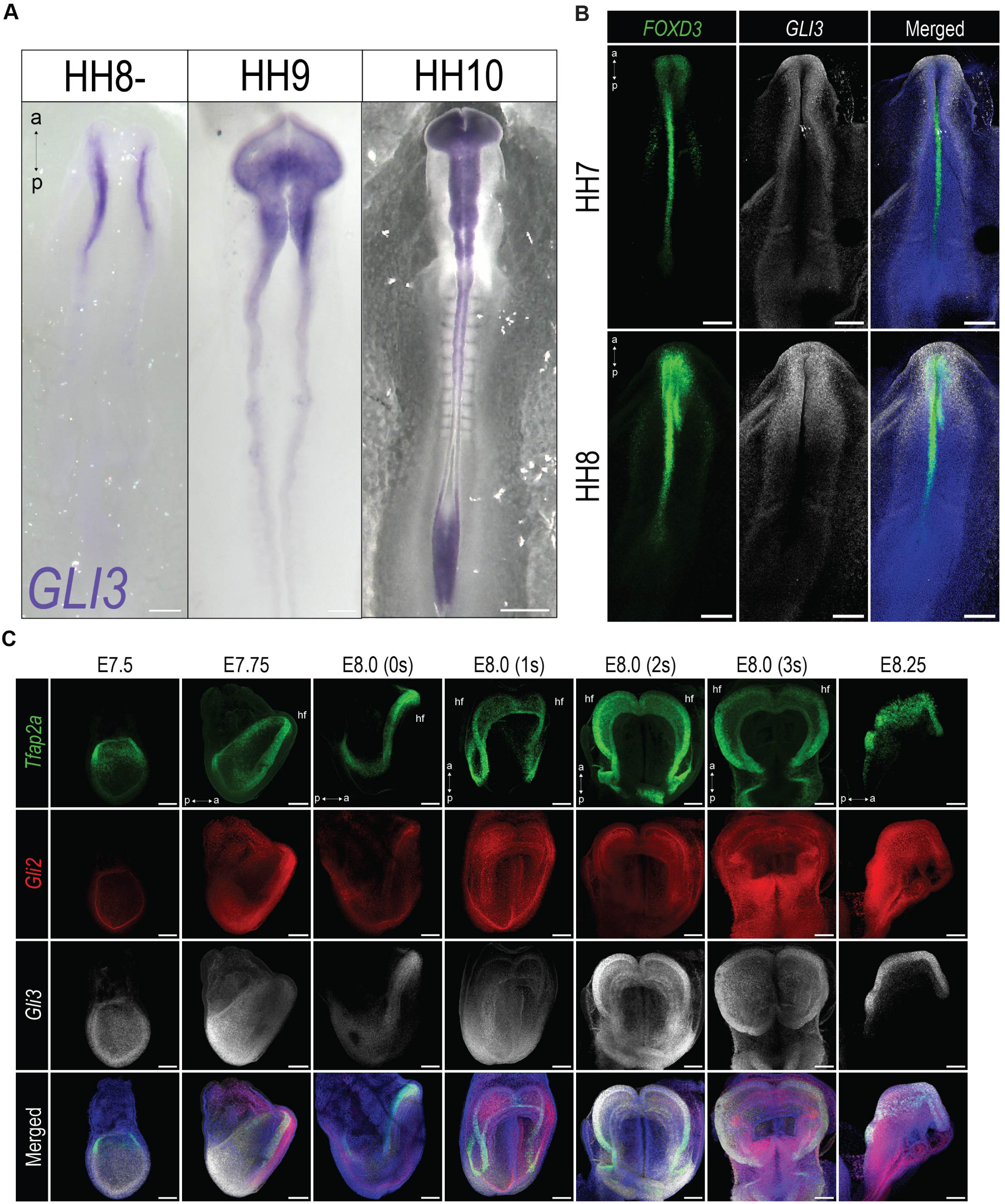
GLI3 expression in developing neural crest cells. (A) *In situ* hybridization for *GLI3* in chick embryos. (B) Whole-mount RNAscope for *FOXD3* and *GLI3* in chick embryos. (C) Whole-mount RNAscope for *Tfap2a*, *Gli2*, and *Gli3* from E7.5-8.25. (hf) headfold. Scale bars: 100μm

**Supplemental Figure S3:**
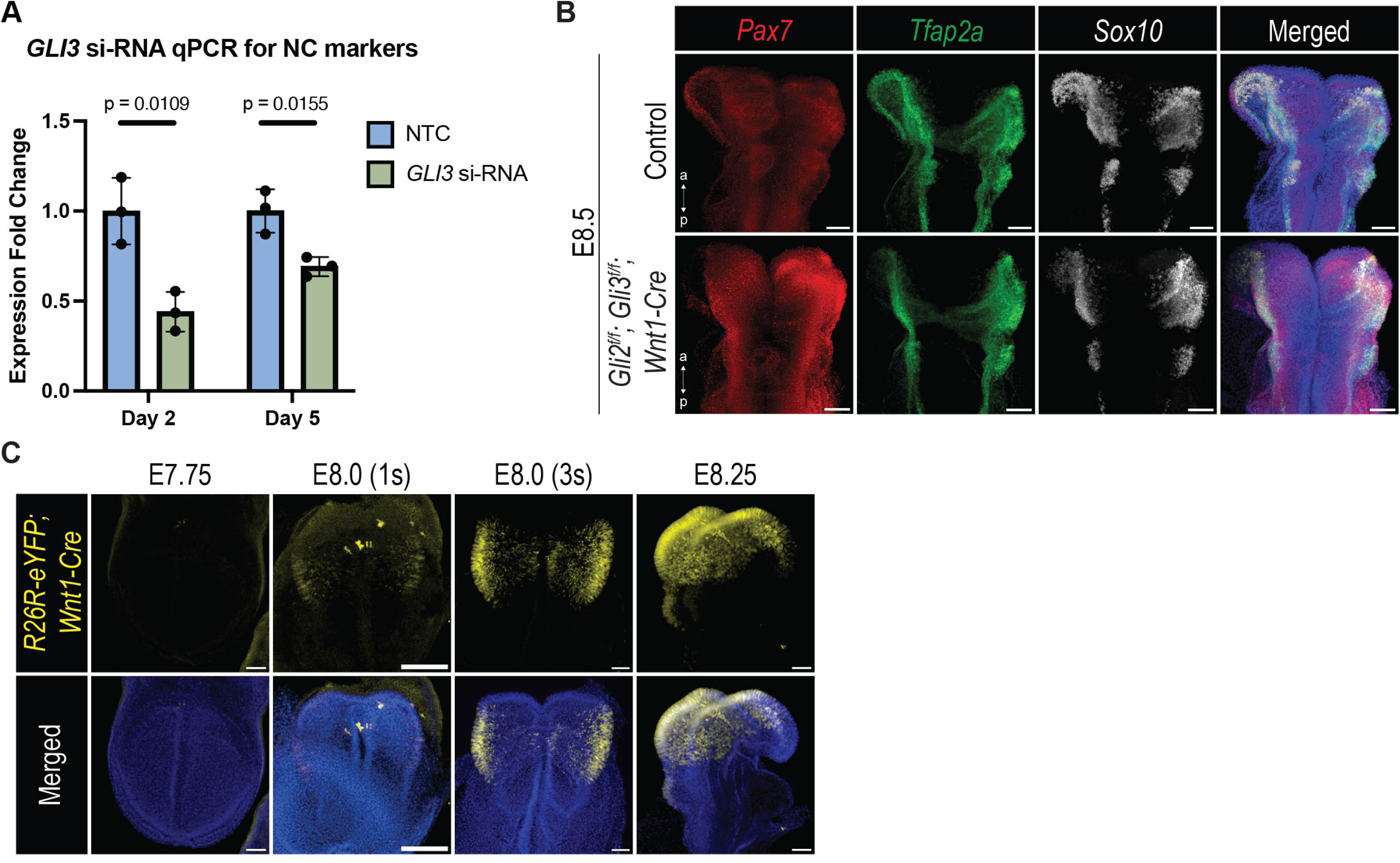
Knockdown of GLI3 prior to neural crest cell specification leads to reduced expression of neural crest cell specifiers. (A) qPCR for *GLI3* to show significant knockdown upon treatment with *GLI3* si-RNA. (B) Whole-mount RNAscope of E8.5 control and *Gli2^f/f^; Gli3^f/f^; Wnt1-Cre* embryos showing maintained expression of neural crest specification genes *Pax7, Tfap2a,* and *Sox10*. (C) Temporal and spatial recombination pattern of *Wnt1-Cre* using *R26R-eYFP; Wnt1-Cre* embryos from E7.75 to E8.25. Scale bars: 100μm.

**Supplemental Figure S4:**
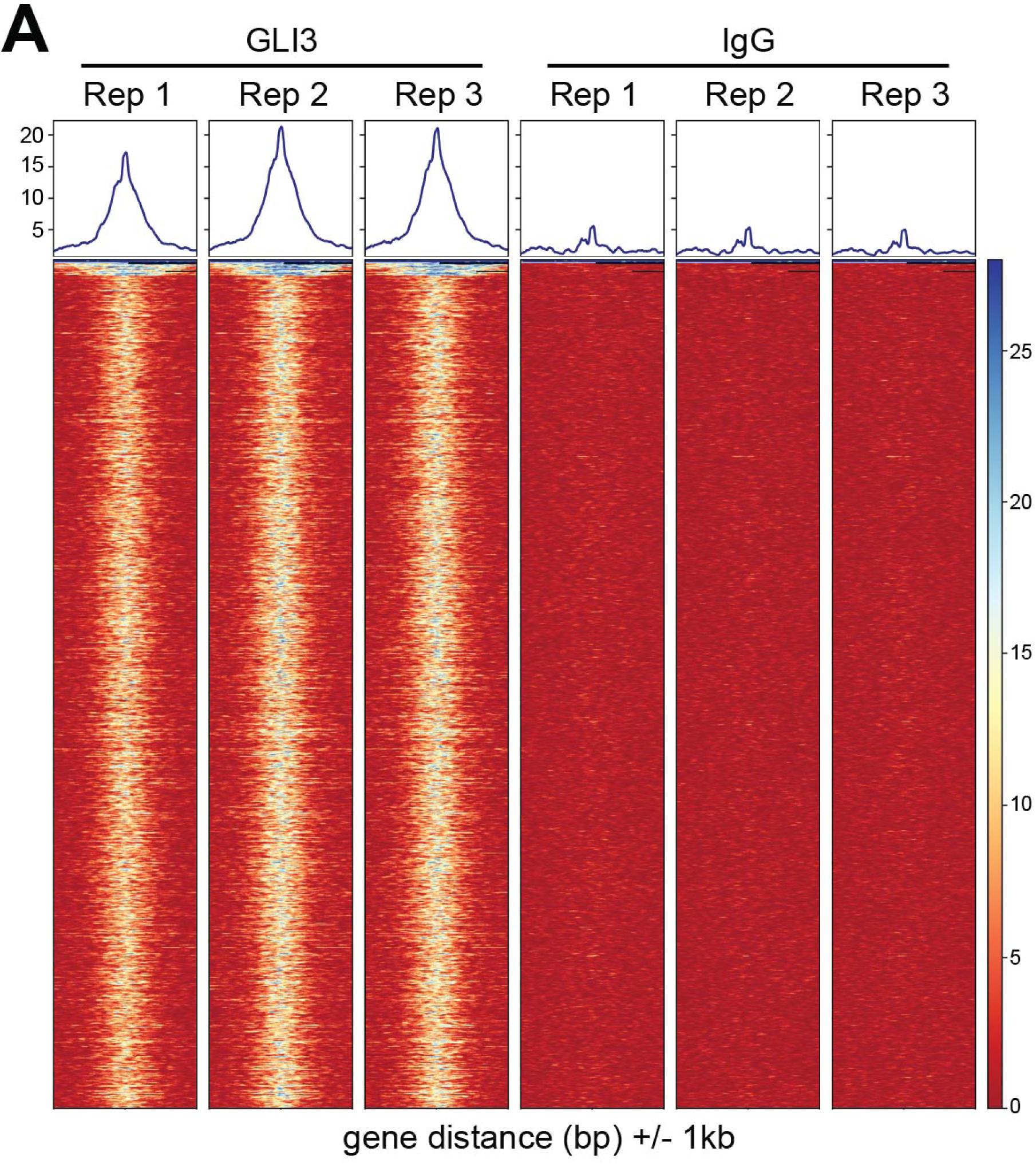
Replicate data for GLI3 CUT&RUN. (A) Heatmap of replicates of data shown in Figure 4B.

**Supplemental Figure S5:**
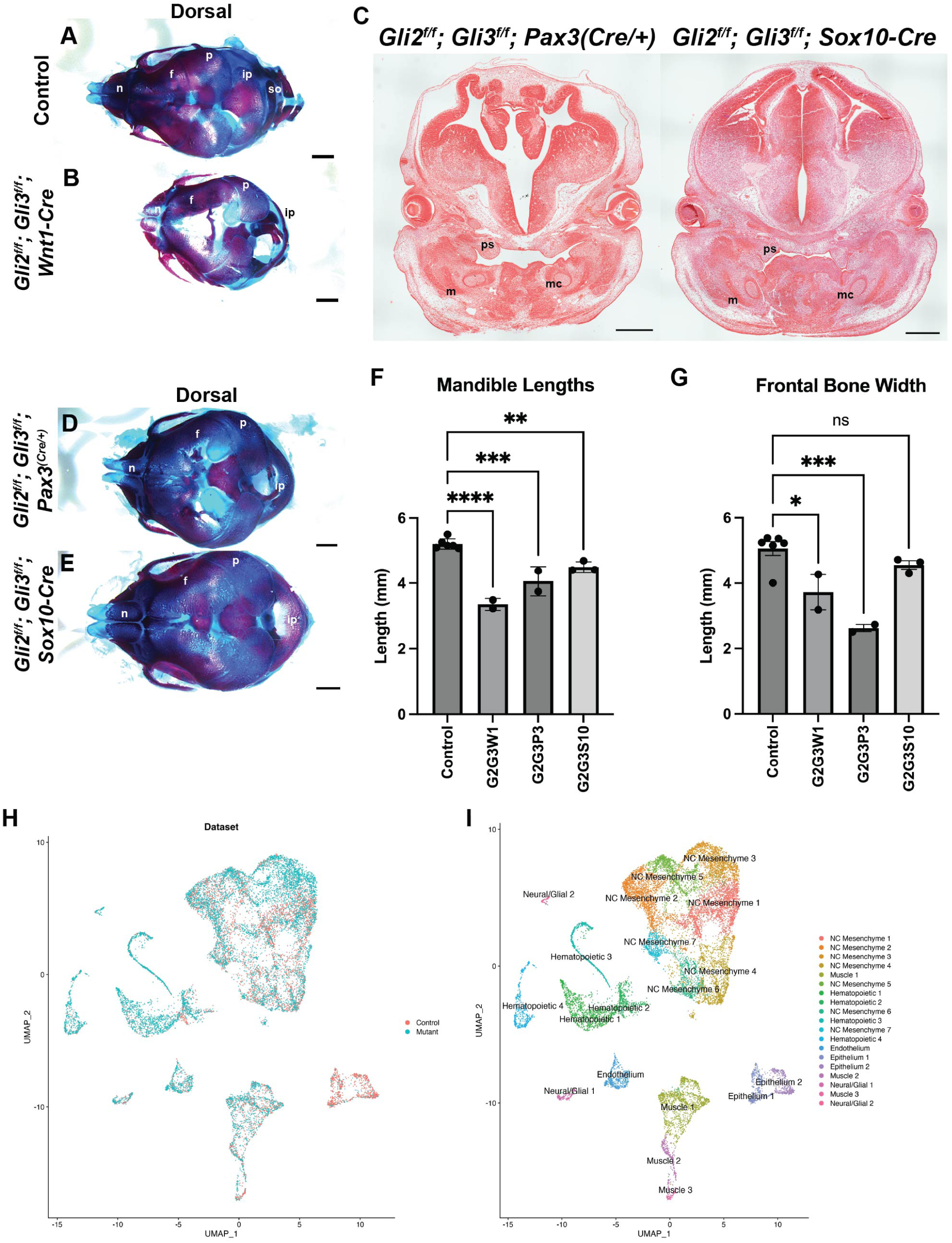
Phenotypes associated with *Gli2^f/f^; Gli3^f/f^* conditional knockouts. (A-B) Dorsal views of Alcian Blue and Alizarin Red stained E18.5 embryonic skulls shown in Figure 6C-F. (C) H&E staining on frontal sections of E13.5 *Gli2^f/f^; Gli3^f/f^; Pax3^(Cre/+)^* and *Gli2^f/f^; Gli3^f/f^; Sox10-Cre* embryos. (D-E) Dorsal views of E18.5 Alcian Blue and Alizarin Red stained embryonic skulls shown in Figure 6G-J. (F-G) Measurements of mandible and frontal bone. (H-I) UMAPs of integrated scRNA-seq data from control and mutant (*Gli2^f/f^; Gli3^f/f^; Wnt1-Cre)* mandibular prominences (MNPs) showing (H) control and mutant and (I) individual clusters. Scale bars: 1mm (A, B, D, E), 500µm (C). * Indicates *P* <0.05, ** indicates *P* <0.005, *** indicates *P* <0.0005, **** indicates *P* <0.0001.

**Supplemental Table1: GLI3 CUT&RUN GO terms.** Pink highlight represents ectomesenchymal GO terms. Purple highlight represents non-ectomesenchymal GO terms.

**Supplemental Table2: Primers sequences for in situ hybridization and qPCR.**

